# Domestication drives changes in floral functional traits that impact generalist pollinator visitation

**DOI:** 10.64898/2026.04.04.716478

**Authors:** Kristen K. Brochu De-Luca, Swayamjit Ray, Avehi Singh, Mila Paiva, Kathleen C. Evans, Carolina Grando, Nash E. Turley, Elizabeth Lavanga, Luis O. Duque, Jared G. Ali, Margarita M. López-Uribe

## Abstract

Crop domestication is an evolutionary process that has led to extraordinary plant phenotypic changes in response to artificial selection. The extent to which domestication has altered floral phenotypes and the implications of these changes for plant-pollinator interactions remain unclear. Here, we characterized the floral phenotypes of wild and domesticated species in the genus *Cucurbita* (squash and pumpkins) and quantified their relative attractiveness to generalist and specialist pollinators. Our results show that functional floral traits change with domestication and that some of these changes impact the visitation of generalist pollinators but not specialists. Crops displayed larger flowers with shorter anthers and wider corollas, lower volatile richness, higher sugar content in pollen, and lower sucrose:glucose ratios in nectar. These trait shifts were associated with pollinator behavior in generalist pollinators, which preferentially visited domesticated flowers. We demonstrate that domestication alters functional plant traits and that these changes affect generalist pollinator preference in agricultural settings.

## Introduction

About 40% of the Earth’s land surface has been converted into agricultural habitats, where domesticated plants are the predominant species (Ellis et al., 2010). This change in land cover has made the traits of crop plants central in shaping ecological interactions (Wood et al., 2015). Through the process of crop domestication, humans have selected plants for traits of agronomic value, like early flowering, increased biomass, and larger, less bitter fruits and seeds, collectively known as ‘domestication syndrome traits’ (Meyer et al., 2012). Artificial selection can also alter plant traits that are not of agronomic interest and can inadvertently reshape ecological interactions between plants and their biotic partners (Chen et al., 2015; Whitehead et al., 2017). For instance, selecting for sweeter fruits is often linked to reductions in secondary metabolites, which can decrease resistance to herbivores (Turcotte et al., 2017; Whitehead et al., 2017; Moreira et al., 2018; Egan et al., 2018) and reduce the attractiveness of domesticated plants to natural enemies of crop pests such as parasitoid wasps (Benrey, 2023). However, the effects of crop domestication on floral traits, which mediate communication with pollinators through cues like shape and scent, remain largely unexplored despite the critical role that pollination plays in crop production (Lorenzo-Felipe et al., 2020).

Crop domestication can shape floral phenotypes through pleiotropic effects resulting from selection on fruit traits. For example, crops generally display larger, more numerous flowers than their wild relatives, likely as a byproduct of selecting for larger fruits and higher yield (Sapir, 2009; Glasser et al., 2023). In addition, crop flowers also tend to produce greater amounts of nectar and pollen than wild flowers (Kuriakose et al., 2009; Turcotte et al., 2017; Whitehead et al., 2017; Moreira et al., 2018; Egan et al., 2018). Another expected consequence of domestication is a reduction in trait variation within species, due to genetic bottlenecks driven by selection aiming to produce highly inbred genetic lines (Meyer & Purugganan, 2013). However, the degree of trait variation within lineages varies throughout the domestication process. In the early stages of domestication, populations often show increased phenotypic variation as new traits emerge and are selected for (Iriondo et al., 2018). Nonetheless, the outcomes of artificial selection on floral traits may be constrained by the dependence on mutualistic interactions with pollinators, and thus, it is unclear to what extent domestication will ultimately affect floral trait changes.

Modifications to floral traits can reshape plant-pollinator interactions by influencing both pollinator attraction and the composition of pollinator communities. In natural systems, floral signals (visual, olfactory, or gustatory) are under strong stabilizing selection because they are essential for attracting pollinators (Knauer & Schiestl, 2015; Gervasi & Schiestl, 2017; Paudel et al., 2019). Changes in floral volatiles, in particular, may play a crucial role in reshaping plant-pollinator communities as they influence both species diversity and their relative abundance (Burkle & Runyon, 2019). This influence is reciprocal: pollinator preferences can also drive the evolution of floral volatiles, leading to changes in the scents emitted by flowers (Schiestl & Johnson, 2013; Zu et al., 2020). However, the ecological consequences of human-mediated changes to functional floral traits on pollinator behavior remain poorly understood (Turcotte et al., 2017). While increased floral display (i.e., larger or more numerous flowers) resulting from domestication can enhance pollinator visitation (Erickson et al., 2022), changes in other floral traits, such as volatiles and floral reward quality, could reduce pollinator attraction (Egan et al., 2018). Alternatively, artificial selection could reinforce or embellish floral traits, increasing pollinator attraction, particularly for crops that rely on animal pollination (Turcotte et al., 2017). As a result, artificial selection may result in a mosaic of traits of agronomic interest and pollinator importance that are distinct from wild relatives.

As agroecosystems expand and intensify globally, they create novel contexts characterized by simplified ecological networks and altered mutualistic interactions (Marrero et al., 2017; Brown & Cunningham, 2019). In these simplified systems, feedback loops between plant traits and pollinator communities likely differ from wild systems. The dominance of generalist pollinators in agricultural systems (Kuriakose et al., 2009; Turcotte et al., 2017; Whitehead et al., 2017; Moreira et al., 2018; Egan et al., 2018) suggests that artificial selection may favor traits that attract generalists or that generalist pollinators can better adapt to the modified floral traits of crops. Generalists often share part of their dietary niche with specialists, but they differ in their foraging needs and preferred floral traits. Specialists rely on their ability to discriminate between host and non-host plants, while generalists prioritize assessing resource quality and respond to chemical signals that are shared by diverse plant species (Schiestl & Johnson, 2013). Consequently, generalists and specialists may use different floral cues while foraging (Brandt et al., 2017), leading to antagonistic preferences on floral phenotypes. If the traits selected during domestication favor generalist pollinators (Waser et al., 1996; Day Briggs & Anderson, 2024), this could create strong feedback loops that promote the generalization of crop pollination systems. However, it is unclear whether domestication will have different impacts on the attractiveness of flowers to generalist vs. specialist pollinators.

Here, we use the pollinator-dependent genus Cucurbita (Cucurbitales: Cucurbitaceae) to test the hypotheses that crop domestication shapes floral functional traits and, in turn, pollinator visitation preferences. The genus Cucurbita (squashes, pumpkins, and gourds) comprises ∼18 species, including six lineages resulting from independent domestication events throughout the Americas: C. pepo pepo (central Mexico), C. pepo ovifera (eastern North America), C. argyrosperma (southern Mexico), C. moschata (likely Central America), C. ficifolia (Peru), and C. maxima (northern Argentina) (Kates et al., 2017; Castellanos-Morales et al., 2018; Chomicki et al., 2020). The multiple independent domestication events within this plant genus offer a unique comparative framework to disentangle the effects of artificial selection from shared ancestry in the investigation of how domestication impacts floral trait evolution. Regarding their pollination system, Cucurbita plants are monoecious with unisexual flowers and thus require pollinators for reproduction (Enríquez et al., 2015; Delgado-Carrillo et al., 2018; McGrady et al., 2020). However, C. foetidissima exhibits gynodioecious populations with some individuals only producing female flowers (Kohn & Biardi, 1995). A highly specialized group of bees known as squash bees (Apidae: Xenoglossa: subgenera Peponapis and Xenoglossa (Freitas et al., 2023)) exclusively collect pollen from these plants (Hurd et al., 1971). In eastern North America, where these crops were introduced 7 kya (Smith & Yarnell, 2009), Cucurbita plants are pollinated by these specialist bees but also several generalist pollinators that forage on a wide variety of host plants, including the Western honey bee (Apis mellifera), several bumble bee species (Bombus spp.), long-horned bees (Melissodes spp.), sweat bees (family Halictidae), and cuckoo bees (Triepeolus spp.) (Artz & Nault, 2011; McGrady et al., 2020). The pollinator communities in the native and extended ranges of the plants comprise both specialist and generalist pollinators (Delgado-Carrillo et al., 2018; McGrady et al., 2020). While bumble bees and honey bees are abundant and effective pollinators in northern latitudes, these bees exclusively collect nectar from Cucurbita flowers and pollinate flowers through passive pollen movement (Artz & Nault, 2011) because consuming their pollen has significant negative effects on their fitness (Brochu et al., 2020).

To investigate the role of domestication on floral traits, we focus on four functional categories that are known to impact pollinator visitation: morphology, floral volatiles, and nutritional reward of pollen and nectar. First, we use a multivariate trait-based approach to test differences in trait space and variation in each trait category from six domesticated and seven wild lineages (Figure 1A). Second, we incorporate a phylogenetic comparative framework to investigate shifts in functional floral traits while controlling for the phylogenetic non-independence among lineages. To assess the role of crop traits on pollinator visitation, we quantified the visitation frequencies of specialist (Xenoglossa pruinosa) and generalist (Bombus spp.) pollinators for a subset of seven wild and domesticated Cucurbita lineages through a randomized block design field experiment over two years. Our combined results empirically demonstrate that domestication has modified different aspects of functional floral traits, and that these changes overall had weak impacts on pollinator preferences. However, the traits modified with domestication were associated with the largest impacts on generalist pollinator visitation, not specialists. Altogether, our results suggest that crop domestication exerts complex selective pressures on floral phenotypes that drive shifts that overall favor generalist crop pollinator systems.

## Materials and Methods

### Greenhouse Experiment

We obtained seeds from 13 Cucurbita lineages from several sources (Table S7). We included one cultivar from five out of the six independent origins of domestication in the genus. The ancestral domestication events occurred in different places of the Americas, but these crops are cultivated across the globe today, where they interact with generalist pollinators (Kates, 2019). Despite the large variation in phenotypes among cultivars within each lineage (e.g., C. pepo), phenotypes of cultivars within a lineage are expected to vary less than cultivars among lineages (Lemoine et al., 2023). We germinated seeds by soaking them in water for 24-48 hours before being transferred to blotting paper. Then, we incubated the seeds at 25°C under a 16:8 hr light:dark cycle until the emergence of hypocotyls to then be transferred to 15 L (four-gallon) pots filled with Metromix in a greenhouse at 28°C using a 16:8 hr light:dark cycle (light intensity at ∼800 umol/s) and fertilized every two weeks with 2 tablespoons of Osmocote 18-5-8 (Griffin greenhouse, USA). We grew plants from November 2017 to April 2018, with sample collection occurring after December 2017. We collected pollen and nectar samples in 1.5 mL centrifuge tubes and stored them at -80°C until used in further analyses. We collected all plant measurements from male flowers due to the larger number of male flowers that plants produce. This focus allowed us to detect potential signs of shifts in traits important for pollinator attraction to these unisexual flowers (Rosas-Guerrero et al., 2011; Sletvold, 2019).

### Morphological Trait Data Collection

We cut flowers directly below the calyx and imaged from 7-10 am with a Canon EOS Rebel T6i from three different perspectives: a top view, a side view, and a sectioned side view to view the anther structures (detailed information for each trait measured can be found in Figures S5-S7). Each photo included a 15 cm ruler that we used to scale the measurements for each flower (Figures S5-S7). To quantify morphological traits, we used the image analysis software ImageJ version 1.21 by measuring 120 flowers (average 12, SD 4.47; Table S8) across 4 wild and 6 domesticated lineages.

### Volatile Trait Data Collection

We measured floral volatiles from 245 different male flowers in total across 6 wild taxa and 7 domesticated lineages, (average 19.5, SD 11.25; ; Table S8). To collect volatiles, we bagged flowers in 12x15 cm oven bags. We collected the volatiles emitted from 6 am to 12 pm in glass filter traps containing HayeSepQ (Sigma Aldrich, USA) by pulling a constant vacuum of air through the filter trap at 250 ml/min and pushing the equivalent amount of charcoal-purified air into the bag. We also used this method to collect control samples from an empty bag each day. Volatiles were eluted from the filter with 150 DL of dichloromethane, and we added 400 ng of nonyl acetate as the internal standard. We analyzed the eluted samples on an Agilent 6890 Gas Chromatograph and 5977 Mass Spectrometer with an Rx-1ms column (30m length, 0.25mm thickness; Restek, USA) by injecting 1ul of sample in splitless mode. We maintained the oven temperature at 40°C for 2 min, then increased the temperature in 10°C increments until it reached 190°C, and then again in 12°C increments until it reached 280°C. Using Chemstation, we identified the compounds in each sample by comparing them to the NIST14 mass spectral library using a cutoff of > 90% fidelity and quantified each compound using the nonyl acetate standard as a reference. The retention indices of all compounds are available in the Supplemental Material (Table S9).

### Nutritional Trait Data Collection

We measured the nutritional characteristics of both pollen (average 4.07, SD 1.8; Table S8) and nectar (average 4.3, SD 2.06; Table S8) from more than 50 flowers across 6 wild taxa and 7 domesticated taxa. We created pooled pollen samples by combining ∼3 mg of pollen from each of the three flowers on a single plant for a total of 9-10 mg of pollen per plant. Pooled pollen samples from at least 3 plants per species were used to conduct a Bradford assay to assess pollen protein content and an assay modified from Van Handel and Day (Day & Van Handel, 1988; Vaudo et al., 2020) to assess pollen lipid and carbohydrate content. Nectar analyses were conducted on a per-flower basis with three technical replicates for each biological sample. We used 2-5ul of nectar to assess both sucrose and glucose using a Roche Sucrose/Glucose enzymatic assay (ID#10139041035). All nutritional assays used at least 3 biological replicates and were quantified with a SpectraMax iD3 Multi-Mode Microplate Reader (Molecular Devices, San Jose, CA, United States). From these values, we also calculated the protein:lipid and sucrose:glucose ratios, which we analyzed in addition to the raw macronutrient values. Our lipid values were lower than those published in other studies, so we suspect some procedural differences led to this discrepancy; however, the relative differences are comparable within our study.

### Pollinator Foraging Experiment

In 2018 and 2019, we conducted field observation experiments (see Figure S8 and S9 for plot design), combining visual assessments with video recordings. We chose three mesophytic wild + domesticated Cucurbita sister species pairs (C. maxima ssp. andreana + C. maxima spp. maxima, C. argyrospermia spp. sororia + C. argyrospermia ssp. argyrosperma, and C. pepo spp. ovifera var. texana + C. pepo ssp. ovifera var. ovifera) in addition to the wild species C. foetidissima to compare bee visitation rates (See Table S6 for full taxa names). Note that although C. pepo ssp. ovifera var. texana is a wild lineage, it is likely derived from C. pepo spp. ovifera var. ovifera (i.e., feral as per (Whitaker & Bemis, 1964; Nee, 1990; Castellanos-Morales et al., 2018) and is considered a domesticated lineage for the analysis. For the pollinator visitation experiments, we placed plants of each species (10 in 2018 and 5 in 2019; Figures S8 and S9) in a randomized row for each of three blocks in a field at the Penn State Agronomy Research Station in Rock Springs, Pennsylvania, USA (40°42’38.1’’ N, 77°57’52.07’’ W), where plots received no insecticide or fungicide applications during the duration of the study, and weed control was conducted mechanically. The surrounding landscape includes a mosaic of research fields, fallow plots, and pasture, within native forest dominated by oaks along with maples, hickories, and associated species on slopes and in woodlots adjacent to agricultural land. We used one field in 2018 and three fields in 2019 (at least 1.35 km from each other). We first grew plants in the greenhouse in May of each year and transplanted them to the field on June 6, 2018, and June 14, 2019. We conducted pollinator observations for 12 days in 2018 between August 8^th^ and 23^rd^ and for 16 days in 2019 between August 1^st^ and 23^rd^. All observations were conducted between 6 am and 11 am, recording temperature and weather conditions on each sampling day. In each row, we counted all male and female flowers and observed floral visitors in each block for 1 minute per plant (10 minutes in 2018 and 5 minutes in 2019). Because we assessed three fields in 2019, we reduced the number of plants per block to make it possible to assess all fields each day. We recorded observations on bee species, bee sex, bee foraging behavior (approach, nectaring, and pollen collecting), and flower sex visited during our visits.

### Statistical Analysis

All statistics were done using R version 4.3.0. To test for differences in trait variances among wild and domesticated taxa, we quantified trait variance within taxa for morphology, volatiles, nectar, and pollen separately. We did this by fitting a non-metric multidimensional scaling (NMDS) ordination using the ’metaMDS’ function and then calculating the average distance to the centroid for each taxa using the ’betadisper’ function (Oksanen et al., 2013). For some traits, we had to remove species that only had a single plant measured. We tested for differences in within-taxa trait variation by fitting phylogenetically controlled generalized least squares (PGLS) models using the ’gls’ function (Pinheiro et al., 2023). In these models, we used a Brownian Motion correlation matrix made with ‘corBrownian’ function (Paradis & Schliep, 2019) using a phylogeny of the focal taxa from Kates et al. (Kates et al., 2017). Models had domestication status as a predictor and the average distance to the centroid for each species as the response. To test for differences in trait space occupied by wild and domesticated taxa, we first averaged all plant data to get one value per trait per plant taxa. For each trait category, we created a Bray-Curtis dissimilarity matrix and visualized differences in trait compositions with NMDS ordinations. We tested for differences in trait compositions between wild and domesticated taxa by fitting PERMANOVA models with the ’adonis2’ function (Oksanen et al., 2013)

To study the impacts of domestication of individual floral traits we again averaged all traits to get one value per plant taxon. Many of the floral morphology traits were highly correlated, so we simplified the data by fitting a principal component ordination using ’prcomp’ function and extracted scores for PC1 and PC2. The values for PC scores were all inverted to make interpretation easier. PC1 was positively correlated with measures of flower size (larger values of PC1 represent larger flowers), and PC2 is positively correlated with corolla angle and negatively correlated with anther length (larger values of PC2 represent more open flowers and shorter anthers; Figure S1). We tested the impact of domestication of each floral trait with PGLS models using the same methods as described above. To visualize the results of all the traits, we extracted model coefficients and 95% confidence intervals (coefficient standard error x 1.96). To allow comparisons among variables, we scaled them to have a mean of 0 and a standard deviation of 1.

We investigated the relationships between plant domestication and pollinator behavior by testing correlations between trait values measured in the greenhouse and pollinator behavior measured from plants in the field. For both datasets, we averaged data to obtain one value for plant taxa and scaled all variables (mean = 0, SD = 1). We used PGLS models to test if domestication status predicted pollinator behavior variables, and also tested relationships between floral traits and pollinator behavior. We extracted coefficients and 95% confidence intervals to visualize the results.

## Results

### Multivariate Trait Space Changes with Domestication

We found significant differences in overall trait space for wild and domesticated lineages in morphological and volatile traits with domestication status in models explaining 40% and 22% of the variation, respectively (Figure 1B, 1D, Table S1). Pollen and nectar traits in wild and domesticated lineages did not significantly differ in trait space (Figure 1F, 1H, Table S1). However, when testing for within-lineage trait variation by comparing distances to species centroids using phylogenetically controlled generalized least squares (PGLS) models, pollen traits in domesticated lineages showed greater variation compared to wild lineages (standardized coefficient = 1.15, Figure 1G, Table S2). We found no differences in within-lineage trait variation for morphological, volatile, or nectar traits (Figure 1C, E, I, Table S2).

**Figure 1.**
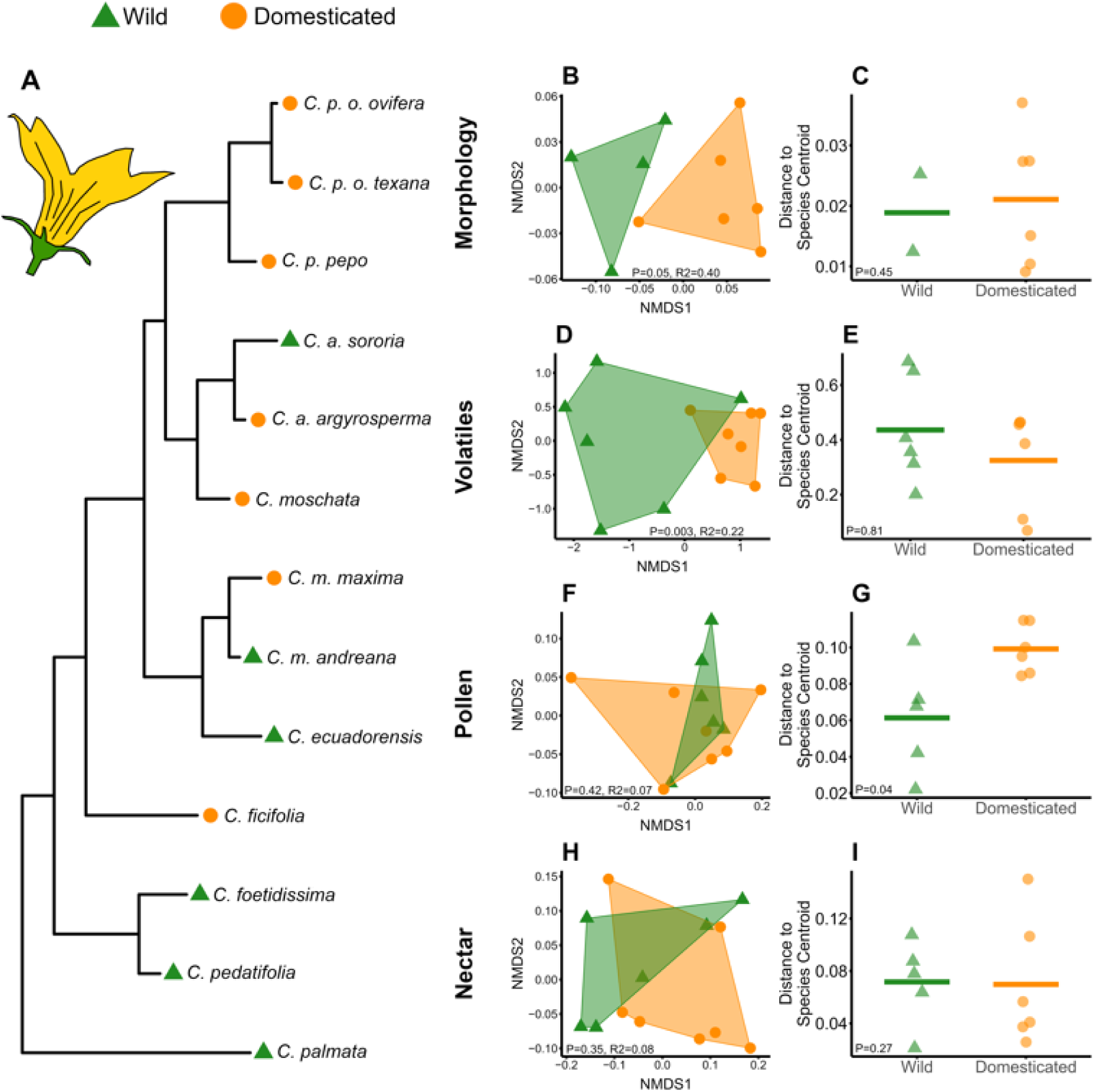
Phylogeny of focal wild and domesticated Cucurbita lineages and effects of domestication on multivariate trait space and within-lineage trait variation. A, Phylogeny of Cucurbita in this study showing multiple independent domestication events. Note that C. pepo ssp. ovifera var texana is a feral lineage, and it is treated as a domesticated species. B, D, F, and H are non-metric multidimensional scaling ordinations showing multivariate trait space, with each point being a species mean for traits measured on overall plant morphology, volatiles, pollen, and nectar, respectively. The statistics on the plots are from a PERMANOVA. C, E, G, and I show variation within lineage using an average distance of individuals within that lineage to the NMDS ordination centroid. P-values are from phylogenetically controlled generalized least squares (PGLS) models. We found significant impacts of domestication on morphology and volatile trait space. Overall, domestication has little effect on within-species variation except for an increase in pollen trait variation.

### Effect of Domestication on Floral Traits

We found that domestication had large effects (standardized coefficients >1) on both PC1 and PC2 (Figure 2, Table S3). PC1 was positively associated with flower size; thus, domesticated plants showed larger flowers overall. PC2 was positively associated with corolla angle and negatively with anther length; therefore, domesticated plants had wider corolla angles and shorter anthers (Figure 2, Table S3). We also found a large effect of domestication on volatile richness (standardized coefficient = - 1.02), with wild plants having greater richness but no difference in volatile abundance (Figure 2, Table S3). For pollen traits, we found that lipids and protein content did not significantly differ between wild and domesticated lineages, but domesticated lineages had higher sugar content (standardized coefficient = 1.03, Figure 2, Table S3). Nectar sucrose and glucose levels did not differ significantly with domestication status, but sucrose:glucose ratio was higher in wild plants (standardized coefficient = -1.3; Figure 2, Table S3).

**Figure 2.**
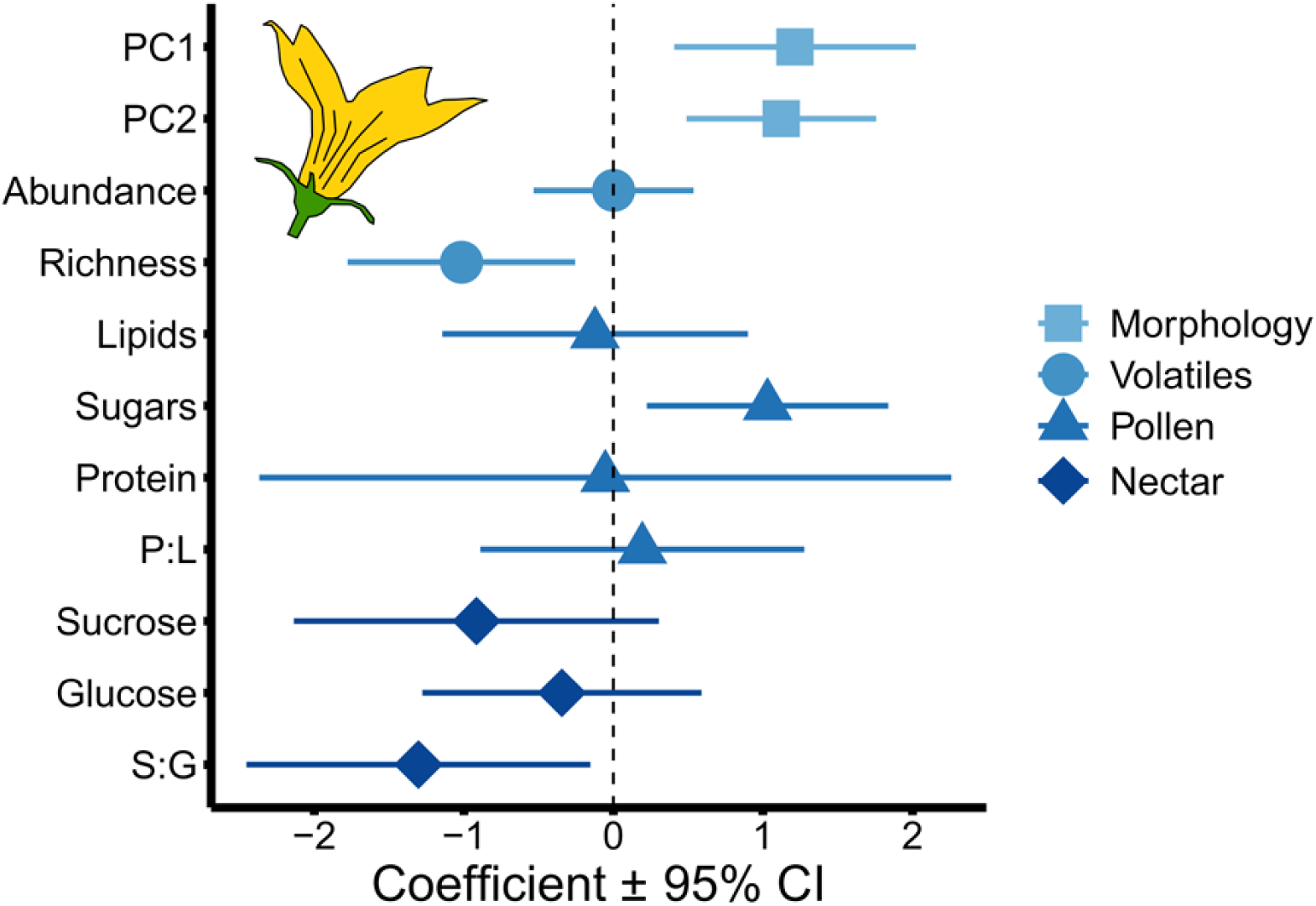
Domestication effects on floral traits. Regression coefficients from phylogenetically controlled generalized least squares (PGLS) models testing the effects of domestication on plant traits, with positive values meaning domesticated plants had larger trait values than wild plants. P:L represents values for pollen:lipid ratios, and S:G represents values for sucrose:glucose ratios. Variables were scaled to have a mean of zero and a standard deviation of 1. Error bars that do not cross zero are significant (P<0.05).

### Effects of Domestication and Floral Traits on Pollinator Visitation

Overall, squash bees (specialists) and bumble bees (generalists) accounted for 98% of the pollinator observations (Table S4). We found trends of generalist pollinators preferring domesticated over wild flowers (Figure 3A, Table S5). Specifically, we found weak evidence that generalist pollinators are more likely to approach (standardized coefficient = 0.8, P = 0.2) and collect nectar (standardized coefficient = 1.2, P = 0.1) on domesticated flowers than wild ones. In contrast, domestication did not impact approaches, nectaring, or pollinating behavior of specialist pollinators (Figure 3A, Table S5). We found evidence for several traits correlating with preference for nectaring by generalist pollinators, but little evidence that Cucurbita flower traits impact visitation in specialist pollinators. For generalist bees, 4 out of 10 floral trait variables had significant correlations with nectaring behavior, while in specialists, none were significant (Figure 3B, Table S6). The strongest response (standardized coefficient = 1.2, P = 0.007) for the generalist pollinator Bombus was to floral morphology PC2, suggesting that Bombus preferred flowers with shorter anthers and more open corollas. The strongest relationship for Xenoglossa was for greater nectaring in plants with increased volatile abundance, but there was only weak evidence for this association (standardized coefficient = 0.8, P = 0.18). Across all floral traits, we found that the traits that had large differences between domesticated and wild species were also the traits that impacted visitation by the generalist pollinator Bombus, but there was no similar relationship in the specialist Xenoglossa (Figure 4). However, with one outlier point removed, Xenoglossa had a similar, though weaker, relationship as Bombus (Figure S2). In the generalist pollinator, there was a positive correlation between the magnitude of floral trait change due to domestication (absolute value of domestication effect size) and the magnitude of the impact (absolute value of effect size) that traits had on Bombus nectaring behavior (Figure 4, t = 3.6, DF = 9, P = 0.006, R^2^ = 0.54). However, that same relationship was not significant when including the directionality of Bombus nectaring effect sizes (Figure S3) because some variables that changed with domestication increased Bombus floral visits (floral morphology, pollen sugars) while others decreased floral visits (volatile richness, nectar sucrose). We also found some evidence for Bombus and Xenoglossa responding differently to specific floral traits; the strongest effect was that increased volatile abundance and richness tended to increase Xenoglossa visitation while decreasing Bombus floral visits (Figure S3, Table S6).

**Figure 3.**
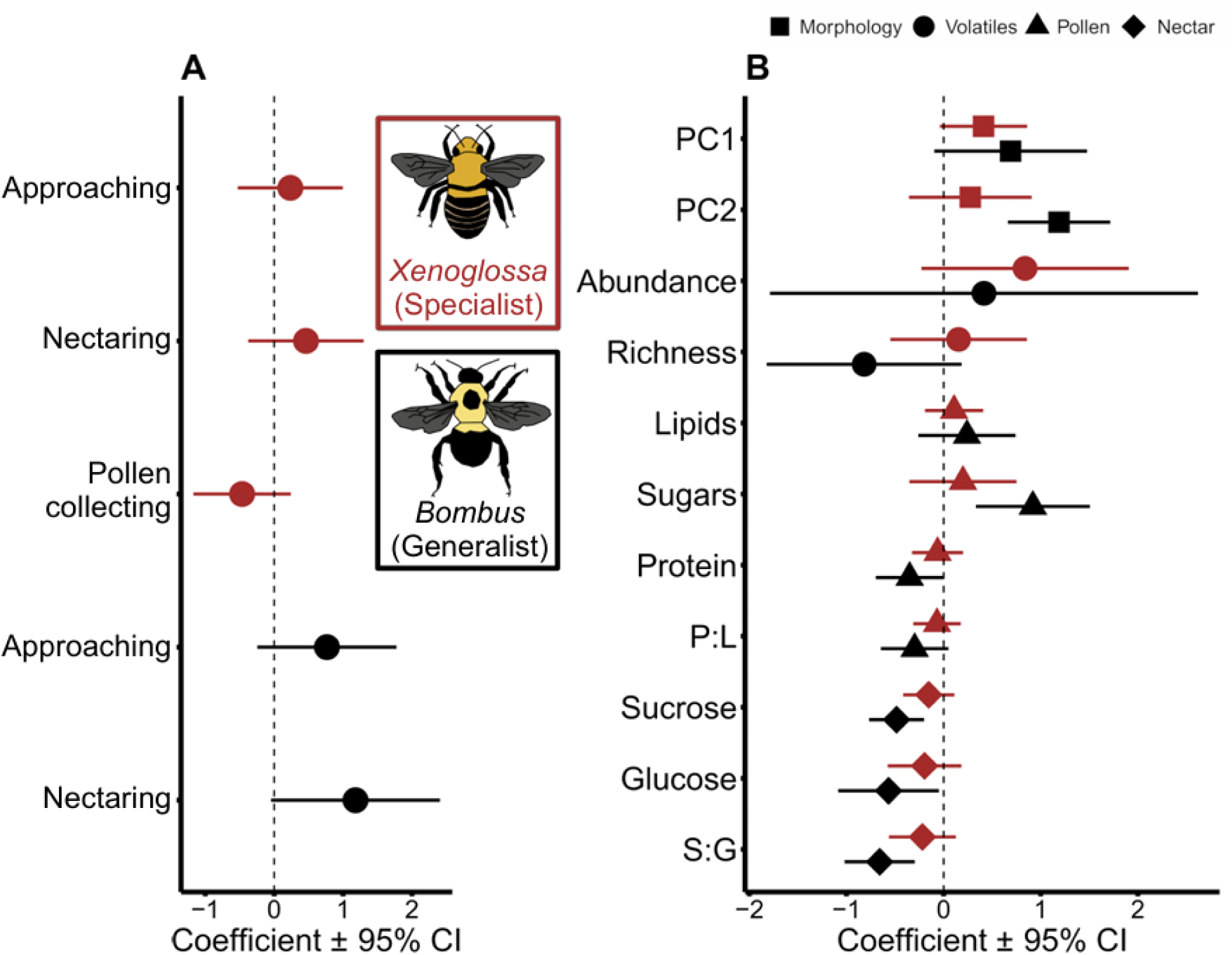
Effects of plant domestication and floral traits on bee pollination behaviors. A. Regression coefficients for models testing the effects of Cucurbita domestication on pollinator behavior of generalist bumble bees (Bombus) and specialist squash bees (Xenoglossa). Positive values indicate that domesticated plants have larger values than wild plants. Models were phylogenetically controlled. B. Standardized coefficients for models testing the impacts of Cucurbita floral traits on pollinator nectaring behavior. Traits are broken down into floral morphology (PC1 related to flower size and PC2 with anther length and corolla angle), volatiles, pollen, and nectar traits. Variables were scaled to have a mean of zero and a standard deviation of 1. Error bars that do not cross zero are significant (P<0.05).

**Figure 4.**
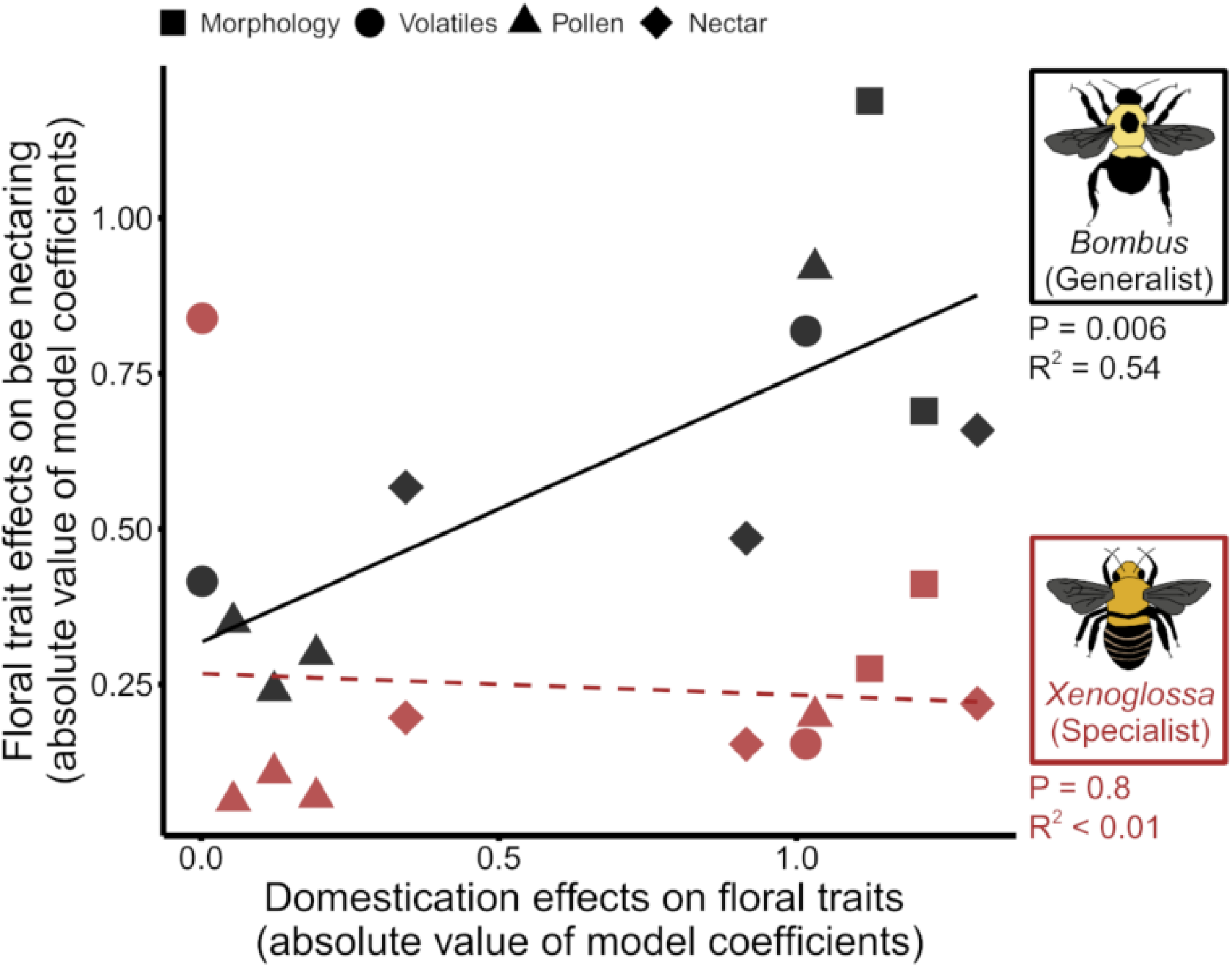
Floral traits changes associated with domestication have greater effects on Bombus nectaring behavior but have no impact on Xenoglossa behavior. Relationship between the magnitude of floral trait change due to domestication (absolute value of regression coefficients from models testing the effects of domestication on plant traits, as shown in Figure 2) and the magnitude of correlation between plant traits and bee nectaring behavior (absolute value of regression coefficients for models testing the impacts of Cucurbita floral traits on pollinator nectaring behavior, as shown in Figure 3B). The statistics above are from linear regression for each bee species.

## Discussion

Overall, our results indicate that domestication alters a variety of floral traits, and those changes have the strongest influence on generalist pollinator behavior. The strongest changes in domesticated flowers were larger flowers with shorter anthers, wider corollas, and reduced volatile richness, as reflected in significant differences in trait space for morphological and volatile traits (Figures 1 & 2; Table S3). These results are consistent with recent studies investigating similar questions about the role of domestication in shaping floral traits in wild and cultivated Cucurbita (Glasser et al., 2023; Villanueva-Espino et al., 2025; Singh et al., 2025). We also observed changes related to higher sugar concentrations in pollen and lower sucrose:glucose ratios in nectar (Figure 2; Table S3). These overall changes in floral traits were positively (morphology and pollen traits) and negatively (volatile and nectar traits) associated with Bombus visitation to flowers (Figure S3). However, domestication overall favored visitations by these generalist pollinators (Figure 3A). These findings suggest that human-mediated changes in floral traits have reshaped interactions with generalist pollinators, which play a dominant role in agroecosystems. In contrast, changes in floral traits were not associated with preferences in the foraging behavior of specialists. This is particularly striking given the assumption that specialist pollinators are more sensitive to floral trait variation (Burger et al., 2013) and poses the possibility of specialist pollinators adapting to novel floral traits. Our findings add to the growing body of literature indicating that crop domestication reshapes the structure of mutualistic interactions, not only for herbivores and root mutualists but also for pollinators (Milla et al., 2015).

Different selective pressures are likely driving phenotypic changes in flowers within this system. For instance, larger inflorescences are a common feature of the domestication syndrome observed in several crops (Ross-Ibarra et al., 2007; Kuriakose et al., 2009; Turcotte et al., 2017; Whitehead et al., 2017; Moreira et al., 2018; Egan et al., 2018), and it implies that the observed increase in Cucurbita crop flower size may be the result of artificial selection on fruit traits or selection on genes with pleiotropic effects. These mechanisms could also apply to other morphological traits, such as anther length and corolla width. The observed reductions in anther length and the increase in corolla width may favor visitation by generalist pollinators in this system (Bombus). With reduced anther length and wider corollas, the large-bodied generalist bees may facilitate greater pollen contact, more effective pollen transfer, and increased plant reproductive success [as per the pollinator-precision hypothesis (Stewart et al., 2022)]. In Cucurbita crop pollination systems, Bombus deposit more pollen grains per flower visit than the specialist Xenoglossa, suggesting that they are more effective pollinators and may impose stronger pollinator-mediated selection (Artz & Nault, 2011; Pfister et al., 2017). At a global scale, generalist pollinators are the dominant visitors to crop flowers, and these interactions are likely imposing new selection pressures that shape functional floral traits in agroecosystems (Brown & Cunningham, 2019).

Another observed pattern was the contrasting behavioral responses of generalist and specialist pollinators to floral trait variation (Figure 4, Table S3). Generalist pollinators more frequently visited flowers with shorter anther lengths and wider corollas, while specialist pollinators showed the opposite trend (Figure S4A). Similarly, lower floral volatile richness was associated with more frequent visits by generalists, whereas the opposite was observed in specialists (Figure S4C). These findings highlight the potential for antagonistic floral preferences between generalist and specialist pollinators. However, the potential for antagonistic responses to traits associated with domesticated flowers is not limited to different pollinators. Herbivores often rely on the same volatile compounds as pollinators to locate their host plants (Theis & Adler, 2012). This preference could result in artificial selection against the production of certain volatile compounds in order to minimize herbivore pressure on crops (Andrews et al., 2007; Theis & Adler, 2012; Turcotte et al., 2017; Whitehead et al., 2017; Moreira et al., 2018; Egan et al., 2018). Losses in the volatile profile of domesticated flowers may, in turn, exert strong selective pressures on the sensory systems of specialists if the compounds that they use to identify flowers are lost during domestication (Singh et al., 2025). Indeed, in its recently expanded range in Eastern North America, X. pruinosa has exclusively foraged on domesticated Cucurbita for thousands of generations (López-Uribe et al., 2016; Pope et al., 2023), and genes related to chemosensation are under strong selection in this region (Pope et al., 2023). This study system may exemplify a case of eco-evolutionary feedback loops resulting from domestication and agricultural environments driving changes in wild pollinators (Turcotte et al., 2017).

Traits related to the sugar content in nectar and pollen were also impacted by domestication, but to a lesser extent, and were associated with contrasting associations in pollinator behavior. We observed no overall differences in pollen protein:lipid ratios between wild and domesticated plants (Figure 2) consistent with other studies in Cucurbita (Villanueva-Espino et al., 2025). The conservation of these key traits for bee foraging preferences and nutritional needs (Vaudo et al., 2016) indicates that these traits are not targets of artificial selection and may be under stabilizing selection. We detected an increase in the number of sugars in pollen, which is critical for pollen fertility (Liu et al., 2021). Interestingly, this increased sugar content was positively associated with Bombus visitation, even though these generalist pollinators neither collect nor consume (Brochu et al., 2020). For nectar traits, we found no consistent changes in the amount of sucrose and glucose. However, the nectar of domesticated flowers showed lower sucrose:glucose ratios than the nectar of wild plants (Figure 3), a trait associated with lower visitation by bees (Figure 3B).

An important consideration in interpreting the results of our study is that the differences we observed between wild and domesticated flowers could have been the result of both artificial (i.e., crop breeding) as well pollinator-mediated selection. While our results cannot distinguish which selective forces drive the observed phenotypic changes, disentangling the role of these complex selective forces would advance our understanding of the causes of plant trait changes during domestication, and potentially their ecological and evolutionary consequences. If floral trait change was driven entirely by artificial selection, then future breeding efforts could select for trait changes to increase pollination services in crop plants. However, if trait changes were mainly driven by contemporary pollinator selection, then new phenotypes in crops are likely already adapted to best maximize yield in response to the pollinator communities in agroecosystems. It is also possible that artificial and pollinators can synergistically act and shape crop floral traits in agroecosystems. Additionally, while we examined floral traits in isolation, many floral traits are perceived by pollinators as integrated phenotypes, meaning that selection pressures may be acting on these traits as units of covarying traits (Junker & Parachnowitsch 2015; Schiestl 2015; Schlichting 1989). Further, our study focused on male floral traits, which may be the subject of weaker selection traits than female flowers that produce fruits and seeds (Villanueva-Espino et al. 2025). Thus, further studies on trait covariation and experimental evolution isolating artificial and pollinator selection on male and female flowers may shed some light on the relative contribution of these selective forces on crop floral phenotypes (e.g., Glasser et al. 2023).

Our results show that crop domestication was associated with shifts in a variety of floral traits, with broader ecological effects that impact pollinator visitation behavior. However, these impacts were dependent on the trait measured and the pollinator; the generalist pollinator was strongly affected by some traits of domesticated flowers, while the specialist was not. While specialist pollinators are typically more sensitive to environmental changes (Bartomeus et al., 2013; Graham et al., 2024), one potential explanation for the lack of responses, in this case, is that they have adapted to changes in floral phenotypes, as the specialist pollinators in this region have predominantly relied on crop flowers for the past thousand years (Pope et al., 2023). Future studies in the ancestral range of the wild plant and specialist pollinator can test this “agricultural tolerance” hypothesis (Brown & Cunningham, 2019; López-Uribe et al., 2025), as populations of the specialist pollinator in their native range are expected to be less adapted to agriculture, and thus prefer to visit wild over domesticated flowers. However, recent studies in the native range of Cucurbita do not support this prediction (Glasser et al., 2023). Generalists often persist in agroecosystems (Kuriakose et al., 2009; De Palma et al., 2015), but our results show that they are still impacted by characteristics of crop plants and that manipulations to the functional traits of crops could both impact their ecology across landscapes and their effectiveness in providing pollination services. Because crops are often cultivated in regions outside their native range (Brown & Cunningham, 2019), and pollinator-dependent crops are increasing globally (Aizen et al., 2008), the trait-driven interactions between crop flowers and pollinators are increasingly important in shaping both wild pollinator populations and crop production. A better understanding of how floral traits are altered in domesticated plants in other pollinator-dependent cropping systems (e.g., Solanum, Prunus, Vaccinium), and what impacts those shifts in traits have on pollinators will provide targets to improve crops in ways that could bolster food security by increasing the attractiveness and utility of flowers to a diversity of wild pollinators.

## ACKNOWLEDGMENTS

This research was supported by a National Science Foundation CAREER Award (DEB-2046474); the USDA NIFA Appropriations under Projects PEN 5010; and the Pennsylvania State University Lorenzo L. Langstroth Endowment funds awarded to MML-U. We thank Scott Diloreto for his help and support in the greenhouse research and Austin Kirk for his assistance with the field experiment. We thank Paul De Luca for his support and help with the field experiment. We also thank members of the López-Uribe, Jha, and Muth labs, Paul De Luca, and Betty Benrey for critical feedback on earlier versions of this manuscript.

## Notes

http://datadryad.org/share/qrutnxUNVQyqQ3MbPLGXk4-3_DZ4KaXu9_r8zNov8D0

### Competing Interest Statement

The authors have declared no competing interest.

