## SupplementalMaterials for "Domestication drives changes in floral functional traits that impact generalist pollinator visitation"

**Domestication MS Supplementary Material**

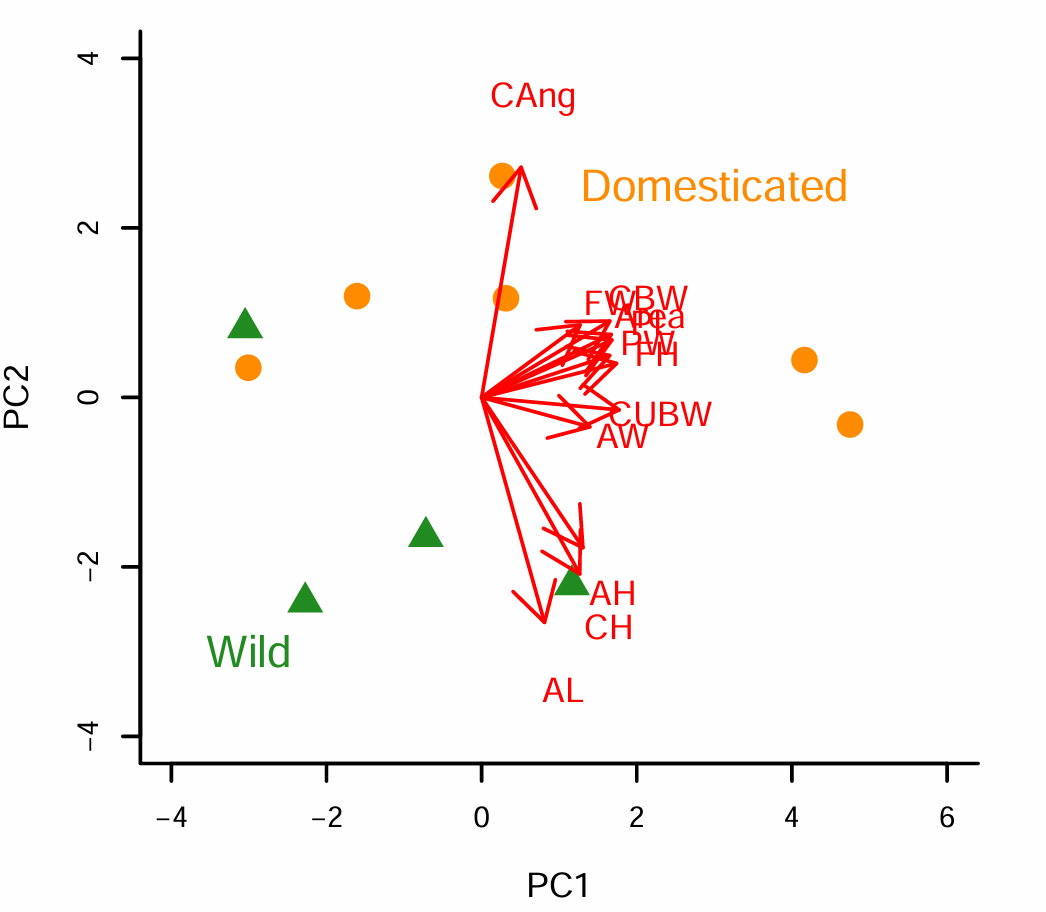

**Figure S1.** Principle components ordination of floral morphology traits of wild and domesticated *Cucurbita* species. Wild species are represented as green triangles and domesticated species as orange circles. The arrows are vectors showing how floral morphology traits relate to ordination space. PC1 is mostly associated with flower size, with larger values (to the right) associated with larger flowers. The two arrows pointing to the right and slightly down are CUBW (corolla upper base width) and AW (anther width), and the nearby cluster of arrows pointing up and to the right are: Area (total floral area from above), CBW (corolla base height), FW (flower height, FW (flower width), PL (petal length), PW (pedal width). PC2 is mostly associated with flower shape and anther size with larger values (up) being wider flowers (more open and less tubular) with smaller anthers. The arrow pointing up is CAng (corolla angle), and the three pointing down are CH (corolla height), and AL (anther length), AH (anther height). Therefore, overall domesticated plants were associated with larger and more open (less tubular flowers) while wild plants were smaller, more tubular, with larger anthers. Note all the values in the PCA were inverted (multiplied by -1) to make interpretations more intuitive. See Figure S5-7 for illustrations of how the floral variables were qualified.

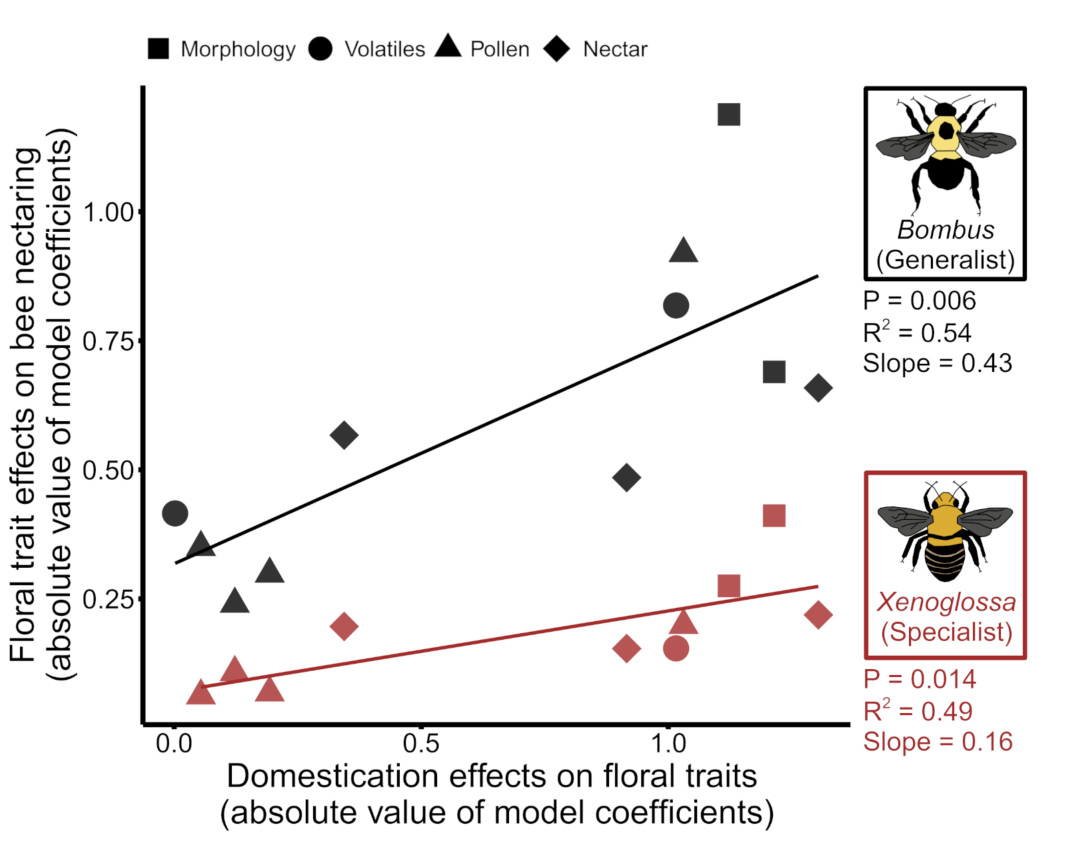

**Figure S2**. Alternative version of Figure 4 from the main text but with an outlier in the *Xenoglossa* data removed. With the volatile abundance outlier data point removed floral traits affected by domestication impact both *Bombus* and *Xenoglossa* nectaring behavior but the strength of the relationship (slope) is 2.7 times larger in *Bombus.* Relationship between the magnitude of floral trait change due to domestication (absolute value of regression coefficients from models testing the effects of domestication on plant traits, as shown in Figure 2) and the magnitude of correlation between plant traits and bee nectaring behavior (absolute value of regression coefficients for models testing the impacts of *Cucurbita* floral traits on pollinator nectaring behavior, as shown in Figure 3B). The statistics above are from linear regression for each bee species.

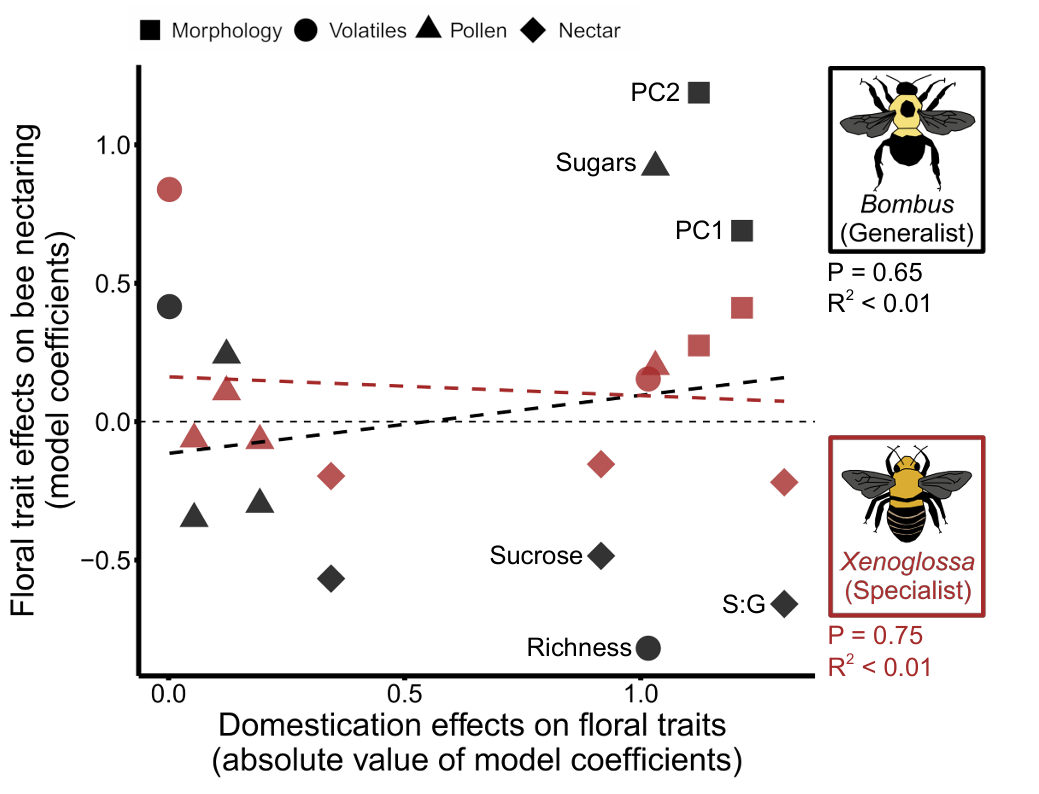

**Figure S3.** Magnitude of floral trait change has no overall relationship with impact that trait has of floral visitation for either *Bombus* or *Xenoglossa*. However, traits which show large differences between wild and domesticated lineages have either strong positive and negative impacts on *Bombus* nectaring. Plot shows the relationship between the magnitude of floral trait change due to domestication (absolute value of regression coefficients from models testing the effects of domestication on plant traits, as shown in Figure 2 and Table S3) and the the correlation between plants traits and bee nectaring behavior (regression coefficients for models testing the impacts of *Cucurbita* floral traits on pollinator nectaring behavior, as shown in Figure 3B and Table S5). Statistics above are from linear regression for each bee species.

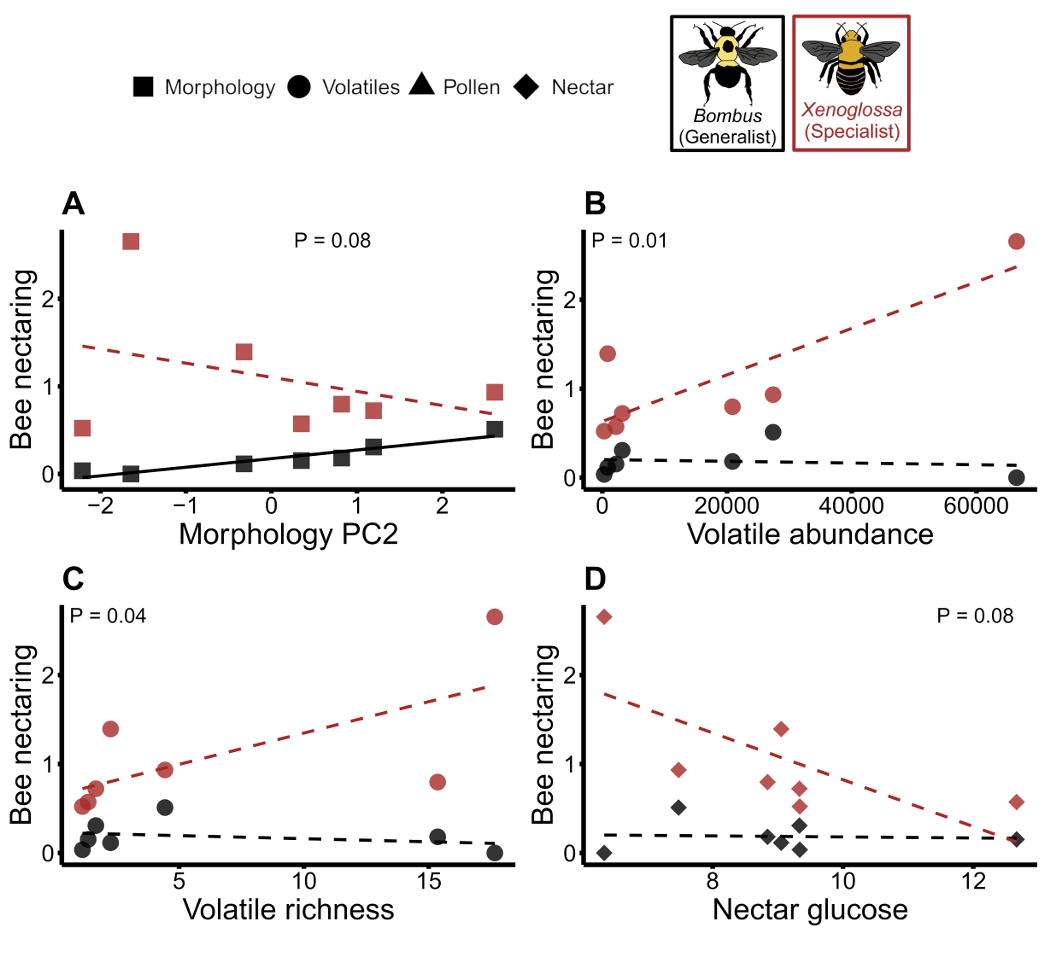

**Figure S4.** Differing impacts of floral traits across 7 *Cucurbita* lineages on nectaring behavior (frequency of floral visits) of both *Bombus* and *Xenoglossa.* P-values on the plot are for plant trait x bee species interaction term in phylogenetically controlled models (see Table S5). Dashed trend lines indicate relationships that were not significant (P<0.05) in models testing each bee species on its own (Table S5).

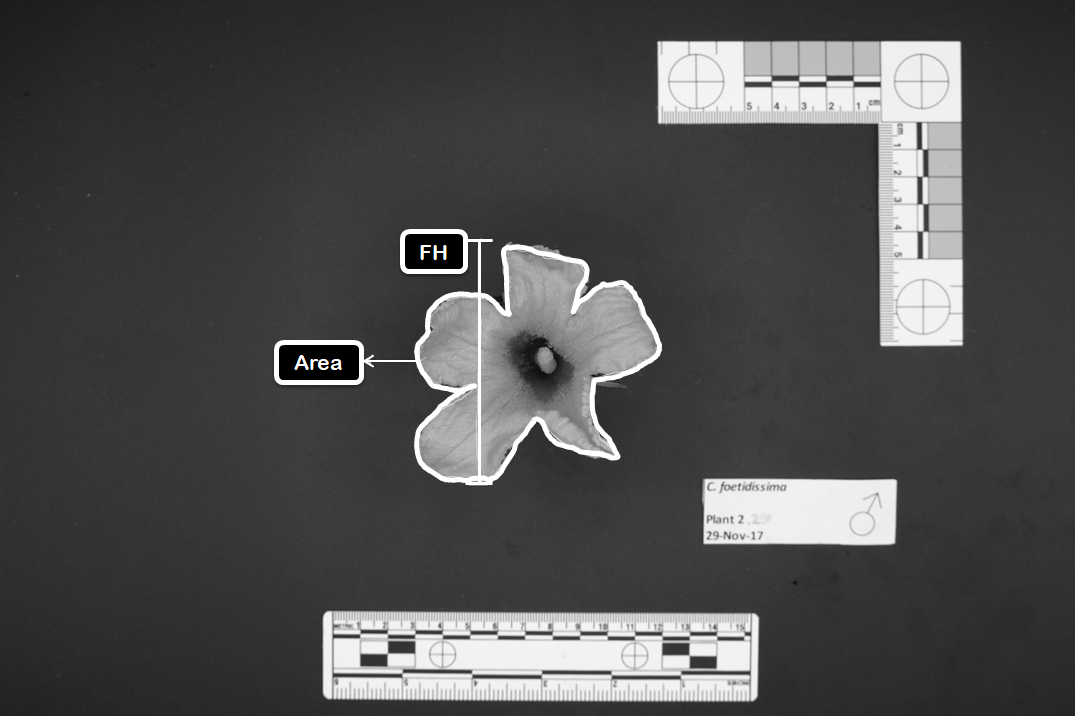
**Figure S5.** Top view of flowers used for morphological measurements indicating Area (Area: the area of the top view calculated using pixel intensities) and Flower Height (FH: the vertical straight line from the highest to the lowest bounds of the flower).

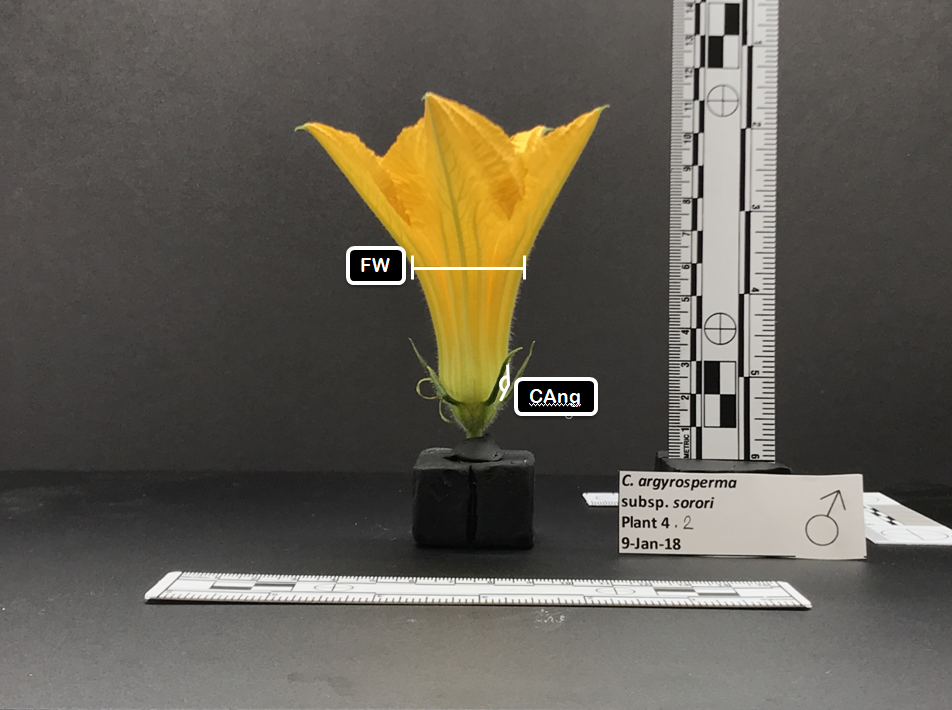

**Figure S6.** Side view of flowers used for morphological measurements indicating Corolla Angle (CAng: the approximate angle made by the corolla as it opens measured in relation to the central axis of the flower) and Flower Width (FW: the width of the flower taken approximately in the middle).

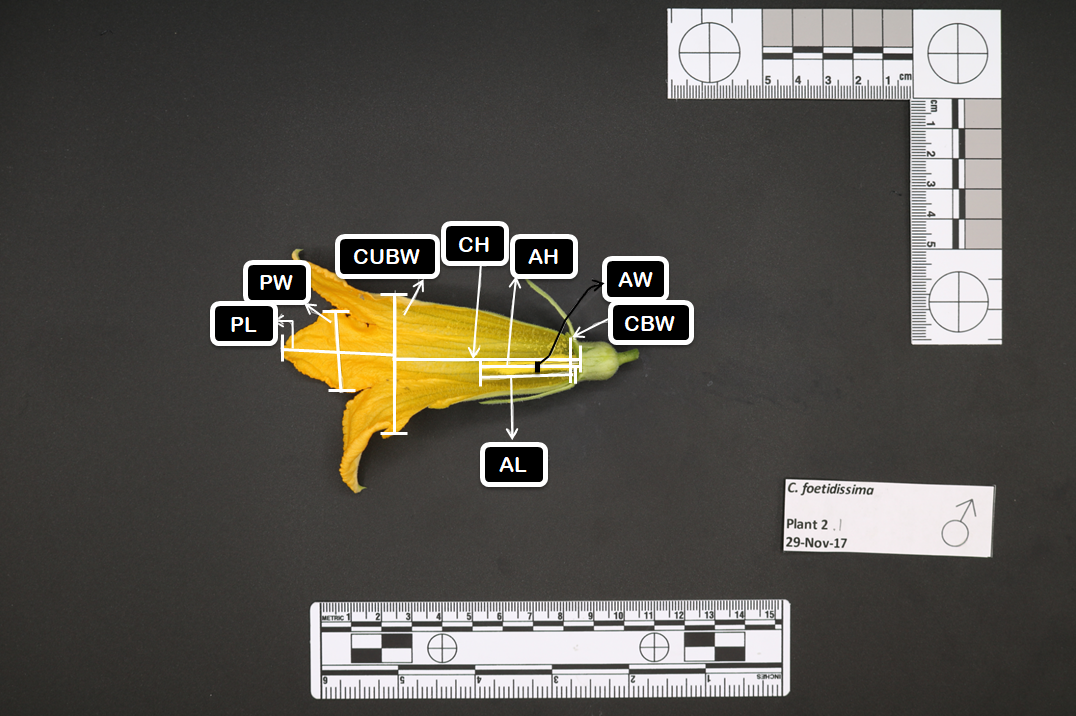

**Figure S7.** Sectioned side view of flowers used for morphological measurements indicating Anther Height (AH: the length from the top of the anther to the base of the calyx), Anther Length (AL: the length from the top of the anther to the base of the corolla), Anther Width (AW: the width of the anther at the middle), Corolla Base Width (CBW: the width of the base of the corolla, at the calyx), Corolla Height (CH: the height of the corolla, from the base to where the petals begin, along the approximate midline of the flower), Corolla Upper Base Width (CUBW: the width of the upper base of the corolla along the cross-section where the petals curve outwards), Petal Length (PL: the length from the tip of the petal to the top of the corolla), and Petal Width (PW: the width of the petal at its broadest point).

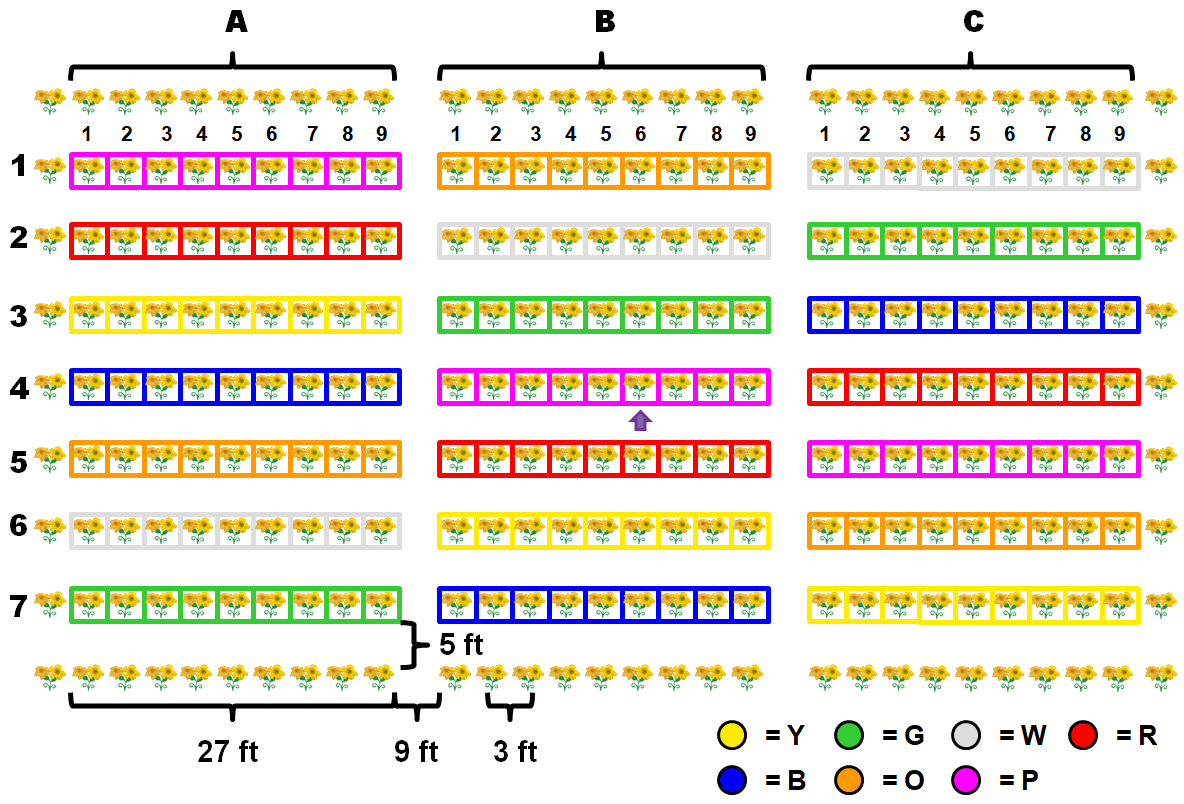

**Figure S8.** Plot design for 2018 field season. Letters represent the three blocks of the experiment, and boxes represent the number of individual plants in each block. Colors represent different species: *C. argyrosperma argyrosperma* (yellow, Y), *C. foetidissima* (green, G), *C. argyrosperma sororia* (white, W), *C. maxima maxima* (red, R), *C. pepo ovifera* var. *texana* (blue, B), *C. pepo ovifera* var. *ovifera* (orange, O), and *C. maxima andreana* (purple, P).

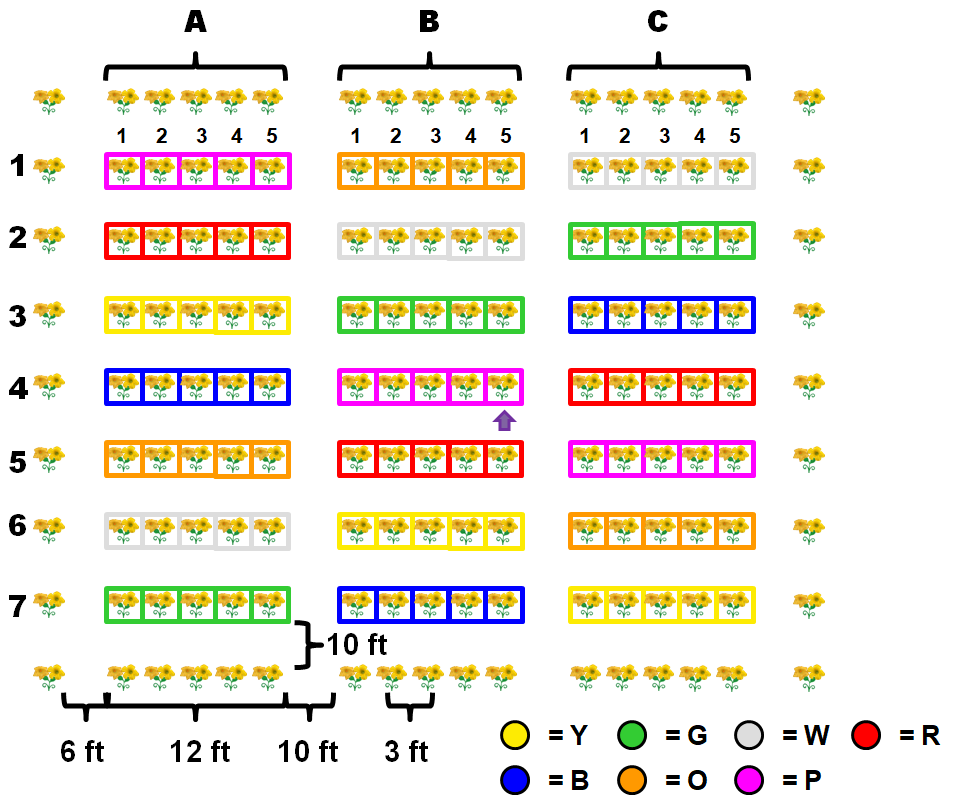

**Figure S9.** Plot design for 2019 field season. This plot design was replicated across three locations (A, B, C). Boxes represent the number of individual plants at each location. Colour represents the *Cucurbita* species planted in that square. Colors represent different species: *C. argyrosperma argyrosperma* (yellow, Y), *C. foetidissima* (green, G), *C. argyrosperma sororia* (white, W), *C. maxima maxima* (red, R), *C. pepo ovifera* var. *texana* (blue, B), *C. pepo ovifera* var. *ovifera* (orange, O), and *C. maxima andreana* (purple, P).

**Table S1.** Results for models testing the effects of plant domestication on multivariate trait space using perMANOVA. All data averaged to have one value per trait per lineage. Bold P-values significant at <0.05.

| **Variable** | **SS** | **F** | **DF** | **P** | **R2** |
| --- | --- | --- | --- | --- | --- |
| Morphology | 0.14 | 5.25 | 1,8 | **0.047** | 0.40 |
| Volatiles | 1.07 | 3.15 | 1,11 | **0.003** | 0.22 |
| Pollen | 0.02 | 0.79 | 1,12 | 0.417 | 0.07 |
| Nectar | 0.03 | 0.90 | 1,11 | 0.351 | 0.08 |

**Table S2**. Results for models testing the effects of plant domestication status on variation within lineages (distance of each individual to lineage centroid on NMDS ordination) using a phylogenetically controlled generalized linear models. All response variables were scaled (mean = 0, SD =1) so that the model coefficients are the predicted changes in standard deviation units in the response variable for going from wild to domesticated lineages, therefore, positive values mean domesticated lineages have a greater predicted value than wild lineages. Bold P-values significant at <0.05.

| **Variable** | **Coef** | **SE** | **t** | **res. DF** | **P** |
| --- | --- | --- | --- | --- | --- |
| Morphology | -1.14 | 1.42 | -0.81 | 6 | 0.450 |
| Volatiles | 0.22 | 0.91 | 0.24 | 10 | 0.813 |
| Pollen | 1.15 | 0.49 | 2.34 | 9 | **0.044** |
| Nectar | -0.81 | 0.70 | -1.16 | 9 | 0.277 |

**Table S3**. Results for models testing the effects of plant domestication status on floral traits using phylogenetically controlled generalized linear models. All response variables were scaled (mean = 0, SD =1) so that the model coefficients are the predicted changes in standard deviation units in the response variable for going from wild to domesticated lineages, therefore, positive values mean domesticated lineages have a greater predicted value than wild lineages. Bold P-values significant at <0.05.

| **Trait** | **Coef** | **SE** | **t** | **res. DF** | **P** |
| --- | --- | --- | --- | --- | --- |
| Morphology PC1 | 1.21 | 0.41 | 2.95 | 8 | **0.019** |
| Morphology PC2 | 1.12 | 0.32 | 3.47 | 8 | **0.008** |
| Volatiles abundance | 0.00 | 0.27 | 0.01 | 11 | 0.995 |
| Volatiles richness | -1.02 | 0.39 | -2.62 | 11 | **0.024** |
| Pollen lipids | -0.12 | 0.52 | -0.24 | 11 | 0.818 |
| Pollen sugars | 1.03 | 0.41 | 2.50 | 11 | **0.029** |
| Pollen protein | -0.05 | 1.18 | -0.05 | 11 | 0.964 |
| Pollen P:L | 0.19 | 0.55 | 0.35 | 11 | 0.733 |
| Nectar sucrose | -0.92 | 0.62 | -1.47 | 11 | 0.169 |
| Nectar glucose | -0.34 | 0.48 | -0.72 | 11 | 0.485 |
| Nectar S:G | -1.30 | 0.59 | -2.22 | 11 | **0.048** |

**Table S4.** Number of floral visitors in the field experiments for the two years of the study. Columns indicate raw numbers and their percentages for each year, and combined across 2018 and 2019.

| **Species Name** | **2018** | | **2019** | | **Total** | |
| --- | --- | --- | --- | --- | --- | --- |
|  | Number | Percent | Number | Percent | Number | Percent |
| *Xenoglossa pruinosa* | 3,941 | 90.45 | 16,282 | 88.01% | 20,223 | 88.6% |
| *Bombus impatiens* | 271 | 6.22% | 2,046 | 11.06% | 2,317 | 10.1% |
| *Apis mellifera* | 114 | 2.62% | 100 | 0.54% | 214 | 0.93% |
| *Melissodes bimaculata* | 2 | 0.05% | 41 | 0.22% | 43 | 0.19% |
| *Triepeolus remigatus* | 0 | 0% | 2 | 0.01% | 2 | 0.009% |
| Halictidae | 29 | 0.67% | 28 | 0.15% | 57 | 0.25% |

**Table S5.** Model results testing effects of domestication on pollinator behavior using phylogenetically controlled generalized linear models. All response variables were scaled (mean = 0, SD =1) so that the model coefficients are the predicted changes in standard deviation units in the response variable for going from wild to domesticated lineages, therefore, positive values mean domesticated lineages have a greater predicted value than wild lineages.

| **Trait** | **Coef** | **res. DF** | **t** | **P** |
| --- | --- | --- | --- | --- |
| *Xenoglossa* approaching | 0.23 | 5 | 0.60 | 0.573 |
| *Xenoglossa* nectaring | 0.46 | 5 | 1.08 | 0.330 |
| *Xenoglossa* pollen collecting | -0.47 | 5 | -1.29 | 0.253 |
| *Bombus* approaching | 0.77 | 5 | 1.49 | 0.197 |
| *Bombus* nectaring | 1.18 | 5 | 1.89 | 0.118 |

**Table S6**. Models testing the relationships between plant floral traits and pollinator behavior using phylogenetically controlled generalized linear models. The first two columns of results are for models with both *Bombus* and *Xenoglossa* data and the table shows just the results for the interaction term between the floral trait and bee species. All response variables were scaled (mean = 0, SD =1) so that the model coefficients are the predicted changes in standard deviation units in the response variable for going from wild to domesticated lineages, therefore, positive values mean domesticated lineages have a greater predicted value than wild lineages. The trait x bee species model coefficients are the differences in slope from going from *Bombus* to *Xenoglossa,* a positive value means *Xenoglossa* had a greater slope than *Bombus.* Interaction models have 10 residual DF and *Bombus* and *Xenoglossa* models have 5 residual DF. Bold P-values significant at <0.05.

|  | **Trait X bee species** | | | | ***Bombus* nectaring** | | | | ***Xenoglossa* nectaring** | | | |
| --- | --- | --- | --- | --- | --- | --- | --- | --- | --- | --- | --- | --- |
| **Variable** | **Coef** | **SE** | **t** | **P** | **Coef** | **SE** | **t** | **P** | **Coef** | **SE** | **t** | **P** |
| Morphology PC1 | 0.40 | 0.35 | 1.12 | 0.290 | 0.69 | 0.40 | 1.72 | 0.147 | 0.41 | 0.23 | 1.80 | 0.133 |
| Morphology PC2 | -0.60 | 0.30 | -1.97 | 0.077 | 1.19 | 0.27 | 4.42 | **0.007** | 0.28 | 0.32 | 0.85 | 0.433 |
| Volatiles abundance | 0.85 | 0.28 | 2.99 | **0.014** | 0.42 | 1.13 | 0.37 | 0.727 | 0.84 | 0.54 | 1.54 | 0.184 |
| Volatiles richness | 0.68 | 0.29 | 2.31 | **0.043** | -0.82 | 0.51 | -1.60 | 0.171 | 0.15 | 0.36 | 0.43 | 0.685 |
| Pollen lipids | 0.08 | 0.35 | 0.23 | 0.823 | 0.24 | 0.26 | 0.94 | 0.391 | 0.11 | 0.15 | 0.70 | 0.514 |
| Pollen sugars | 0.30 | 0.33 | 0.92 | 0.381 | 0.92 | 0.30 | 3.07 | **0.028** | 0.20 | 0.28 | 0.71 | 0.511 |
| Pollen protein | -0.20 | 0.38 | -0.53 | 0.605 | -0.35 | 0.18 | -1.97 | 0.106 | -0.06 | 0.13 | -0.46 | 0.662 |
| Pollen P:L | -0.28 | 0.37 | -0.75 | 0.473 | -0.30 | 0.18 | -1.68 | 0.154 | -0.07 | 0.13 | -0.55 | 0.609 |
| Nectar sucrose | -0.38 | 0.35 | -1.08 | 0.306 | -0.48 | 0.14 | -3.35 | **0.020** | -0.15 | 0.13 | -1.14 | 0.306 |
| Nectar glucose | -0.59 | 0.30 | -1.95 | 0.080 | -0.57 | 0.26 | -2.14 | 0.085 | -0.20 | 0.19 | -1.01 | 0.357 |
| Nectar S:G | -0.02 | 0.38 | -0.05 | 0.962 | -0.66 | 0.18 | -3.58 | **0.016** | -0.22 | 0.18 | -1.24 | 0.269 |

**Table S7**. Sources of seeds for each plant species: Bobby Seeds (<https://www.bobby-seeds.com>); Burpee (<https://www.burpee.com/>); Desert Seed Store (no longer available); GRIN-Global USDA (<https://www.ars-grin.gov/>, <https://www.grin-global.org/>); Johnny’s Selected Seeds (<https://www.johnnyseeds.com>); Magic Garden Seeds (<https://www.magicgardenseeds.com/>); Native Seeds Search (<https://www.nativeseeds.org>); Rare Plants Europe (<http://www.rareplants.es>); Seeds Savers (<https://www.seedsavers.org/>); Strictly Medicinal Seeds (<https://strictlymedicinalseeds.com/>); Trade Winds Fruit (<https://www.tradewindsfruit.com>).

| **Full taxa name** | **Abbreviated taxa names** | **Domestication status** | **Seed sources** |
| --- | --- | --- | --- |
| *C. palmata* | *C. palmata* | Wild | Desert Seed Store (Coyote Melon Gourd) |
| *C. pedatifolia* | *C. pedatifolia* | Wild | GRIN-Global USDA (PI 442341) |
| *C. foetidissima* | *C. foetidissima* | Wild | GRIN-Global USDA; Strictly Medicinal Seeds (Buffalo Gourd) |
| *C. ficifolia* | *C. ficifolia* | Domesticated | Thomas Andres; Magic Garden Seeds (Fig-Leaved Gourd); Trade Winds Fruit (Fig-Leafed Gourd) |
| *C. ecuadorensis* | *C. ecuadorensis* | Wild | Thomas Andres; GRIN-Global USDA (PI 432442 02) |
| *C. maxima* ssp. *andreana* | *C. m.* *andreana* | Wild | Thomas Andres; GRIN-Global USDA (PI 458653); Bobby Seeds (*Cucurbita andreana*); Rare Plants Europe (Wild Giant Squash) |
| *C. maxima* ssp. *maxima* | *C. m.* *maxima* | Domesticated | GRIN-Global USDA; Seed Savers (Golden Hubbard Squash) |
| *C. moschata* | *C. moschata* | Domesticated | Johnny’s Selected Seeds (Waltham Butternut) |
| *C. argyrosperma* ssp. *argyrosperma* | *C. a. argyrosperma* | Domesticated | GRIN-Global USDA; Native Seeds Search (EA003) |
| *C. argyrosperma* ssp. *sororia* | *C. a. sororia* | Wild | Native Seeds Search (EA040) |
| *C. pepo ssp.pepo* | *C. p.* *pepo* | Domesticated | Johnny’s Selected Seeds (Yellow Crookneck); Native Seeds Search (EP050) |
| *C. pepo ssp. ovifera var. texana* | *C. p. o.* *texana* | Domesticated | Andy Stephenson; GRIN-Global USDA |
| *C. pepo ssp. ovifera var. ovifera* | *C. p. o. ovifera* | Domesticated | GRIN-Global USDA; Burpee (Tennessee Dancing Gourd) |

**Table S8**. Number of plants and flowers examined for each taxa for each trait category.

| **Taxa** | **Trait category** | **Num. of plants** | **Num. of flowers** |
| --- | --- | --- | --- |
| *C. palmata* | Morphology | 1 | 2 |
| *C. pedatifolia* | Morphology | – | – |
| *C. foetidissima* | Morphology | 4 | 6 |
| *C. ficifolia* | Morphology | – | – |
| *C. ecuadorensis* | Morphology | – | – |
| *C. maxima ssp. andreana* | Morphology | 7 | 13 |
| *C. maxima ssp. maxima* | Morphology | 4 | 14 |
| *C. moschata* | Morphology | 4 | 15 |
| *C. argyrosperma ssp. argyrosperma* | Morphology | 3 | 16 |
| *C. argyrosperma ssp. sororia* | Morphology | 4 | 13 |
| *C. pepo ssp.pepo* | Morphology | 3 | 14 |
| *C. pepo ssp. ovifera var. texana* | Morphology | 7 | 15 |
| *C. pepo ssp. ovifera var. ovifera* | Morphology | 4 | 12 |
| *C. palmata* | Volatiles | 5 | 20 |
| *C. pedatifolia* | Volatiles | 3 | 10 |
| *C. foetidissima* | Volatiles | 5 | 16 |
| *C. ficifolia* | Volatiles | 1 | 4 |
| *C. ecuadorensis* | Volatiles | 4 | 5 |
| *C. maxima ssp. andreana* | Volatiles | 6 | 16 |
| *C. maxima ssp. maxima* | Volatiles | 9 | 29 |
| *C. moschata* | Volatiles | 9 | 28 |
| *C. argyrosperma ssp. argyrosperma* | Volatiles | 6 | 20 |
| *C. argyrosperma ssp. sororia* | Volatiles | 4 | 6 |
| *C. pepo ssp.pepo* | Volatiles | 6 | 21 |
| *C. pepo ssp. ovifera var. texana* | Volatiles | 13 | 47 |
| *C. pepo ssp. ovifera var. ovifera* | Volatiles | 7 | 23 |
| *C. palmata* | Pollen | 2 | 4 |
| *C. pedatifolia* | Pollen | 1 | 3 |
| *C. foetidissima* | Pollen | 4 | 2 |
| *C. ficifolia* | Pollen | 1 | 1 |
| *C. ecuadorensis* | Pollen | 4 | 3 |
| *C. maxima ssp. andreana* | Pollen | 4 | 4 |
| *C. maxima ssp. maxima* | Pollen | 4 | 5 |
| *C. moschata* | Pollen | 4 | 5 |
| *C. argyrosperma ssp. argyrosperma* | Pollen | 4 | 5 |
| *C. argyrosperma ssp. sororia* | Pollen | 4 | 5 |
| *C. pepo ssp.pepo* | Pollen | 4 | 5 |
| *C. pepo ssp. ovifera var. texana* | Pollen | 4 | 6 |
| *C. pepo ssp. ovifera var. ovifera* | Pollen | 4 | 9 |
| *C. palmata* | Nectar | 3 | 4 |
| *C. pedatifolia* | Nectar | 3 | 3 |
| *C. foetidissima* | Nectar | 1 | 2 |
| *C. ficifolia* | Nectar | 1 | 1 |
| *C. ecuadorensis* | Nectar | 3 | 3 |
| *C. maxima ssp. andreana* | Nectar | 4 | 4 |
| *C. maxima ssp. maxima* | Nectar | 4 | 5 |
| *C. moschata* | Nectar | 5 | 5 |
| *C. argyrosperma ssp. argyrosperma* | Nectar | 5 | 5 |
| *C. argyrosperma ssp. sororia* | Nectar | 4 | 5 |
| *C. pepo ssp. pepo* | Nectar | 6 | 5 |
| *C. pepo ssp. ovifera var. texana* | Nectar | 3 | 6 |
| *C. pepo ssp. ovifera var. ovifera* | Nectar | 5 | 9 |

**Table S9**. Retention indices for all volatile compounds. Abbreviations for compound class are as follows:

| **Compound** | **Compound Class** | **Retention Index** |
| --- | --- | --- |
| Myrcene | HC | 989.34 |
| Ocimene | HC | 1035.09 |
| methoxymethyl_benzene | AR | 977.69 |
| trimethylbenzene | AR | 988.24 |
| Benzaldehyde | AR | 937.29 |
| Hexanol_1_ethyl_2 | HC | 1022.78 |
| Limonene | AR | 1029.82 |
| Decanal | HC | 1207.03 |
| Undecane | HC | 1099.93 |
| Dodecane | HC | 1200.07 |
| Linalool | HC | 1093.33 |
| alpha_Funebrene | TER | 1416.24 |
| EDimethylnonatriene_4_8 | HC | 1113.59 |
| Dimethoxybenzene_1_4 | AR | 1146.18 |
| Ethyl_acetophenone | TER | 1261.60 |
| Indole | TER | 1264.35 |
| Trimethoxybenzene_1_2_4 | AR | 1340.21 |
| Phenylethane1_nitro_2 | AR | 1261.60 |
| Bicyclopentyl_2_one | TER | 1269.69 |
| Tridecane | HC | 1308.71 |
| methylCyclopentenone | AR | 1393.08 |
| Dimethylhexahydronaphthalene | TER | 1401.74 |
| E_beta_Farnesene | TER | 1456.58 |
| Tetradecanol | HC | 1484.62 |
| Hexadecanol | HC | 1684.04 |
| Tetradecane | HC | 1408.64 |
| Pentadecane | HC | 1502.45 |
| Hexadecane | HC | 1600.90 |
| Octadecane | HC | 1801.81 |
| Pentadecanol | HC | 1779.86 |
| Cedranone | TER | 1504.76 |
| E_Nerolidol | TER | 1557.68 |
| alpha_Copaene | TER | 1388.09 |
| b_Elemene | TER | 1399.55 |
| GermacreneD | TER | 1440.57 |
| a_Muurolene | TER | 1543.24 |
| E_beta_Ionone | TER | 1336.56 |
| Tetrahydro_trimethhyl_naphthalene | TER | 1323.01 |
| E_Caryophyllene | TER | 1434.47 |
| alpha_Gurjunene | TER | 1643.37 |
| gamma_Muurolene | TER | 1460.38 |
| Amorpha_diene | TER | 1467.19 |
| Italicene | TER | 1483.20 |
| betaCubebene | TER | 1490.02 |
| alpha_Humulene | TER | 1465.69 |
| gamma_Curcumene | TER | 1483.20 |
| trans_a_Bergamotene | TER | 1489.30 |
| alpha_Cedrene | TER | 1427.65 |
| E_a_Farnesene | TER | 1490.81 |
| alpha_Zingiberene | TER | 1495.40 |
| beta_Bisabolene | TER | 1513.61 |
| Tetradecene | HC | 1490.04 |
| Hexadecene | HC | 1583.67 |
| Cyclodecene | HC | 1676.68 |
| Heptadecene | HC | 1583.67 |
| Alpha_ambrinol | TER | 1456.97 |
| Geranyl_acetone | TER | 1436.77 |
| Farnesal | TER | 1734.02 |
| beta_Himachalene | TER | 1520.78 |
| gamma_Cadinene | TER | 1459.59 |
| Butylated_hydroxytoluene | AR | 1504.76 |
| delta_Amorphene | TER | 1527.21 |
| gamma_E_Bisabolene | TER | 1534.39 |
| cis_alpha_Bisabolene | TER | 1541.65 |
